# Adaptation in somatosensory afferents improves rate and temporal coding of vibrotactile stimulus features

**DOI:** 10.1101/2025.05.06.652367

**Authors:** Adel Halawa, Christopher Dedek, Laura Medlock, Dhekra Al-Basha, Stéphanie Ratté, Steven A Prescott

## Abstract

Adaptation is a common neural phenomenon wherein sustained stimulation evokes fewer action potentials (spikes) over time. Rather than simply reduce firing rate, adaptation may help neurons form better (i.e. more discriminable) representations of sensory input. To study effects of adaptation in low-threshold mechanoreceptors (LTMRs), we recorded single unit LTMR responses to 30-second-long vibrotactile stimuli with different intensities and frequencies applied to the hind paw of rats. To assess the impact of adaptation on somatosensory encoding, decoders were applied to the initial and late (adapted) phases of population-level responses to assess the decodability (discrimination) of stimulus intensity and frequency. Adaptation-mediated changes in the rate and timing (phase-locking) of spikes were quantified. Rate coding of stimulus intensity was improved by the nonuniform reduction in firing rate across responses to different stimuli, and across neurons. This improvement was absent in simulations with uniform reductions in firing rate, thus revealing the necessity of stimulus-dependent variability in adaptation effects. Spike timing (quantified as interspike intervals) remained highly informative about stimulus frequency throughout stimulation despite the progressive reduction in spike count over time. When the drop in spike count was accounted for, adaptation was found to improve temporal coding of stimulus frequency by increasing the precision of phase-locking. In other words, adaptation improved the precision of spike timing, and this increased the information about stimulus frequency conveyed by each spike. These results show that adaptation, by modulating spiking in different ways, can improve encoding of different stimulus features using different coding schemes.

**SIGNIFICANCE STATEMENT:** Firing rate adaptation refers to the progressive reduction in spiking during sustained stimulation. Adaptation is often ascribed to fatigue and could conceivably introduce ambiguity into neural representations by causing (the probability or timing of) spike generation to depend not only on the current stimulus, but also on past stimulation. By recording from rat primary somatosensory afferents during vibrotactile stimulation of the paw, we show that nonuniform adaptation across sets of afferents improves rate-based coding of stimulus intensity. We also show that adaptation improves spike timing (phase-locking to a periodic stimulus) which, in turn, improves temporal coding of stimulus frequency. These results demonstrate that early-stage adaptation improves somatosensory signals relayed centrally.

## INTRODUCTION

Codes – the representation of information by a set of abstract symbols – are at the heart of neuroscience, genetics, and much more. Diverse neural coding strategies exist but can be broadly grouped by whether the rate of spikes (counted over a relatively long-time window) or the timing of individual spikes (relative to other spikes in the same neuron, to spikes in other neurons, etc.) carry relevant information (1–3). At the population level, the relative co-activation of differently tuned neurons (4–6) or their relative spike-timings (7–10) can encode information (11). These so-called rate and temporal coding schemes are not mutually exclusive but, rather, enable multiplexed coding of different stimulus features (10, 12–15).

Regardless of the inferred coding strategy, the set of spike trains a neuron can generate in a fixed time is finite and, therefore, its coding capacity is upper bounded (16–18). Capacity is further limited by the neuron’s firing rate distribution and the amount of noise in the system (16, 19). Nevertheless, primary sensory neurons are responsible for encoding all the information about the world to which the brain has access, and they must do so under diverse conditions. How do sensory neurons make the most of their limited information carrying capacity? Adaptive encoding is one solution.

Adaptive encoding involves neurons adjusting their encoding function depending on context (20–26). For instance, the range of light intensities experienced in a darkened theater or while outside on a sunny day differ dramatically, but our retinas manage to discriminate small variations that occur within each context (27). With adaptation, the encoding function can be optimized to maximize mutual information between light intensity and neural responses in each new context (20–30). Adaptive encoding has been rigorously demonstrated throughout the visual pathway – in retina (27, 28), lateral geniculate nucleus (31, 32), V1 (23, 33–35), and higher-order visual regions (36, 37) – and across species – in flies (20–22), salamanders, rabbits (27, 28), mice (34, 35), cats (23, 32, 33), and monkeys (31, 36, 37). In the somatosensory system, human psychophysical studies have shown that adaptation improves discrimination of the intensity and frequency of vibrotactile stimuli (38, 39), but animal studies have focused on central mechanisms (for reviews see 40, 41), leaving it unclear how adaptation in primary afferents contribute. Adaptation at this earliest stage may help or hinder subsequent processing.

To investigate adaptive encoding in somatosensory afferents, we recorded in vivo from low-threshold mechanoreceptors (LTMRs) innervating the glabrous skin of rat hind paws. We stimulated anesthetized rats with 30-second-long vibrotactile stimuli of various intensities and frequencies, and analyzed the decodability of population responses, especially how decodability was affected by LTMR adaptation. Consistent with previous reports (8, 42–48), we found that stimulus intensity is best encoded by population firing rate whereas stimulus frequency is best encoded by spike times. We also found that adaptation improved the decodability of both stimulus features. The basis for improvement differed between coding strategies: nonuniform (i.e. stimulus-dependent) changes in tuning improved the coding efficiency (bits/sec) of the rate code whereas more precise spike-timing improved the metabolic efficiency (bits/spike) of the temporal code.

## RESULTS

### Adaptation effects are not uniform across stimulus parameters nor across LTMRs

LTMR firing rates typically decreased during sustained (30 s-long) stimuli but the degree and even the direction of firing rate changes varied across stimulus parameters. **Figure 1A** illustrates firing rate plotted over time for an LTMR stimulated at various frequencies with a 200 mN force; firing rate remained constant for 20-40 Hz stimuli (based on consistent 1:1 entrainment) but decreased over time for other stimulus frequencies. **Figure 1B** illustrates similar heterogeneity for responses to different stimulus intensities, with responses to some forces decreasing (50, 75, and 100 mN), responses to other forces remaining unchanged (125, 225 mN), and the response to 200 mN stimulation initially increasing.

**Figure 1.**
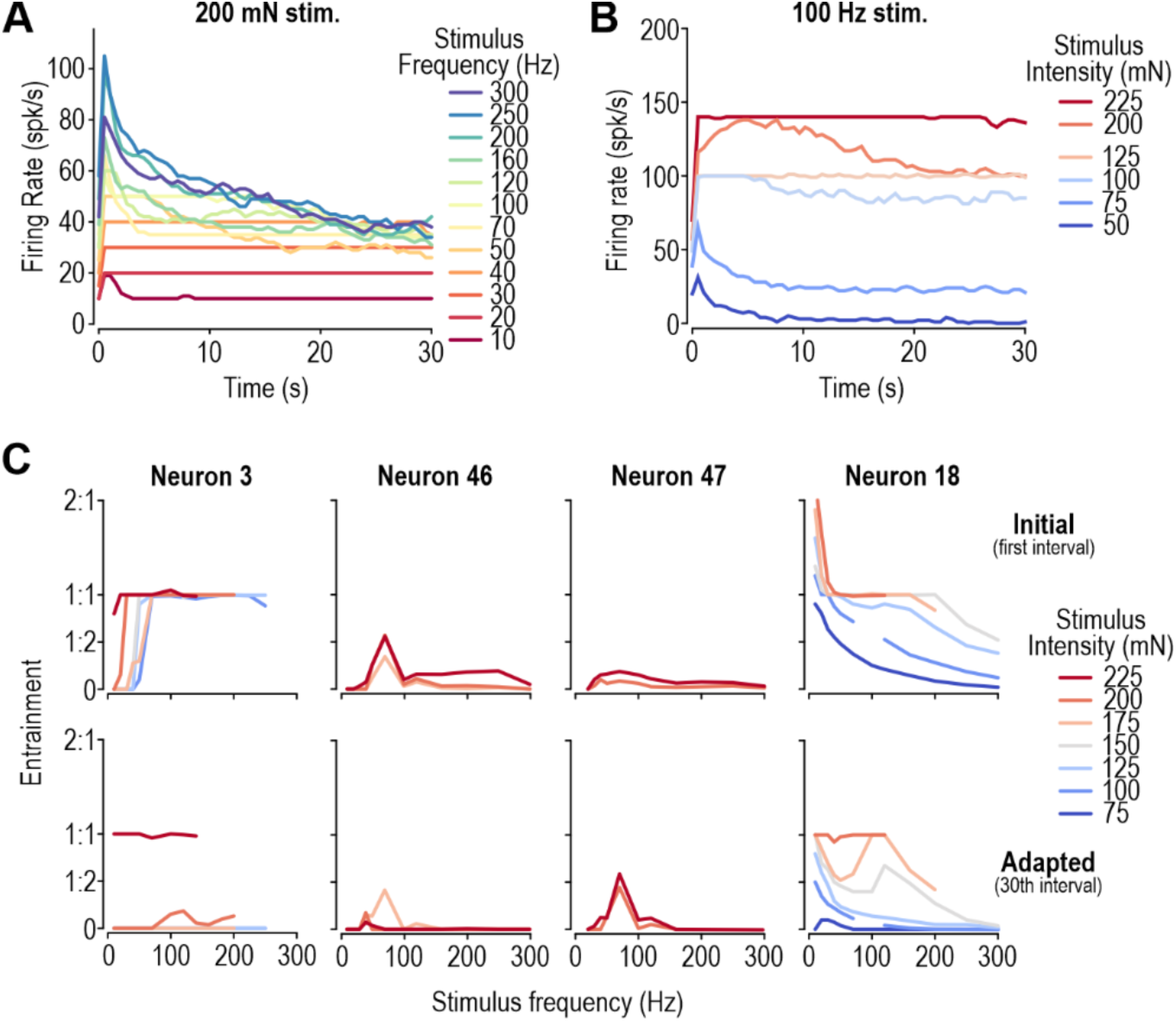
Adaptation effects depend on stimulus parameters and LTMR identity. **A.** Evolution of firing rate over time for sample LTMR stimulated at different vibration frequencies with 200 mN force. Firing rate was calculated in 500 ms-long bins and then smoothed with a 1 s-long sliding window. **B.** Like A, but for responses to 100 Hz stimulation at different intensities. **C.** Tuning curves of four example LTMRs based on firing rates measured during the first (top) and 30th (bottom) second of stimulation. Missing data represent incomplete recordings or unreliable spike sorting. Adapted tuning curves are not uniformly re-scaled versions of initial tuning curves because of differences in adaptation across stimulus parameters (see text).

The sensitivity of adaptation effects to stimulus parameters suggests that tuning curves change nonlinearly over time, implying an LTMR’s adapted tuning curve is not a simple linear re-scaling of its original tuning curve. Comparing tuning curve changes across sample LTMRs reveals the diversity of adaptation effects (**Fig. 1C**). For example, Neuron 3 decreased its entrainment to low-intensity but not high-intensity stimuli, whereas Neuron 46 exhibited the opposite pattern. Changes also depended on stimulus frequency, with adaptation selectively increasing the response of Neuron 47 to 70 Hz stimulation whereas adaptation in Neuron 18 revealed a previously ambiguous preference for 120 Hz stimulation with medium-to-high intensity. This diversity of adaptation effects reveals that adaptation does not reduce firing rate by a fixed amount; instead, the evolution of LTMR responses depends on stimulus parameters and differs from one LTMR to the next.

### Adaptation improves encoding of stimulus intensity by the population firing rate

How do changes in LTMR firing rates affect encoding of vibrotactile stimuli? We addressed this question at the population level since encoding likely involves a distributed code (6, 8, 43–45, 48, 49). One way to monitor how encoding changes is to measure mutual information between the neural response and the stimulus. Since direct calculation of mutual information (22, 50) is intractable with a population code, we opted to measure a lower-bound on mutual information using a decoder (17, 51–53). The firing rate of each neuron was calculated using 200 ms-long bins and a population vector was created from eight LTMRs (**Fig. 2A**). To track adaptation effects, each 30 s-long response was divided into 1 s-long intervals and, for each interval, a different support vector classifier (SVC) was trained and tested on its ability to classify the intensity or frequency of the stimulus using the firing rate vectors (see Methods).

**Figure 2.**
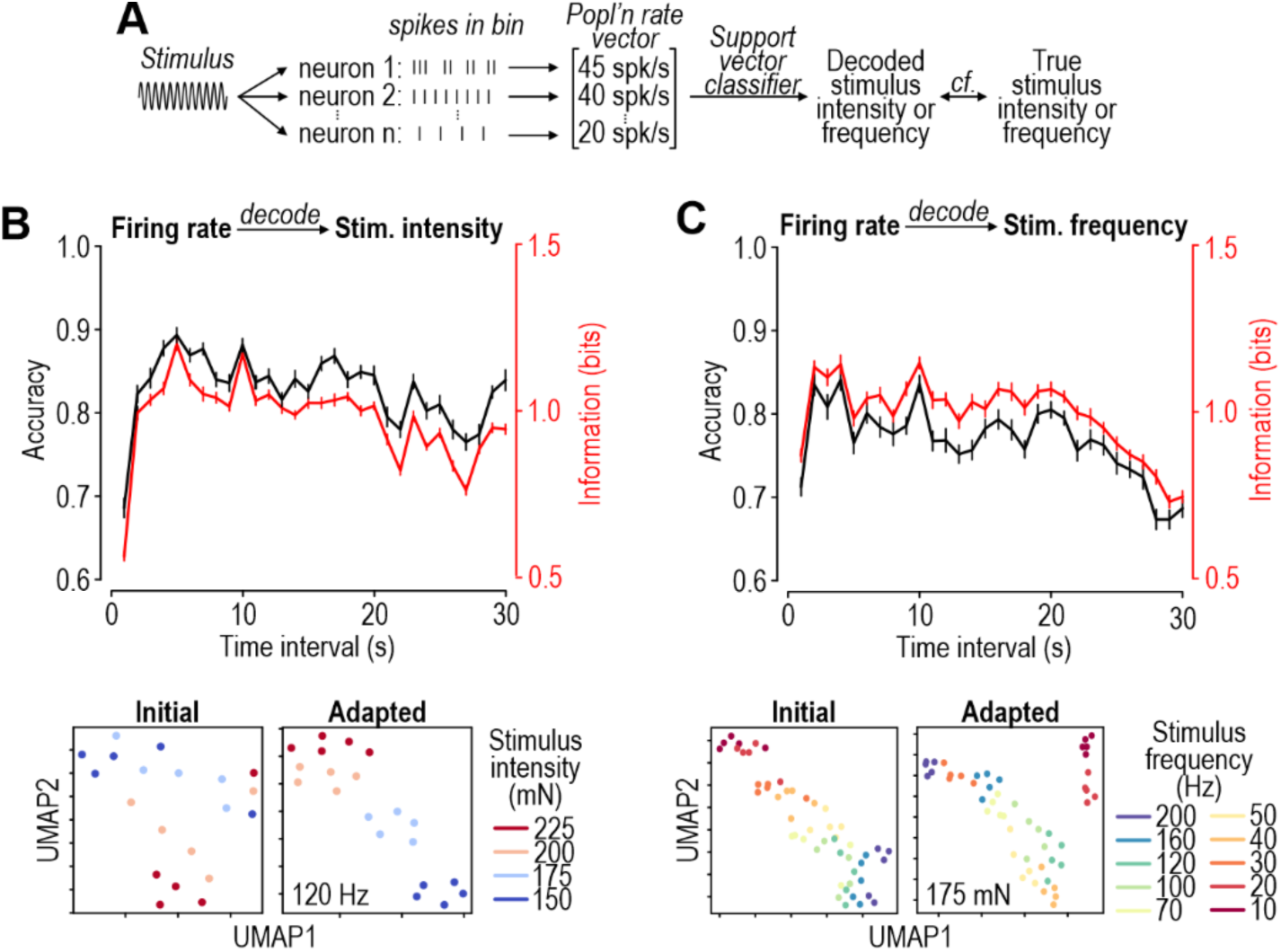
Firing rate adaptation improves population rate coding of stimulus intensity. **A.** Procedure for evaluating population rate coding. Firing rate for each LTMR is determined using bins to create a vector of rates from which we infer stimulus intensity or frequency. The population rate vector is recalculated at 1-sec intervals to compare changes in coding over time. A different classifier was trained and tested for each interval with the train/test split (0.7/0.3) repeated 20 times with different splits. **B.** Decoding of stimulus intensity quantified as decoding accuracy (black) or as lower bound on mutual information (red). Both metrics (mean ± SEM) exhibit a sustained increase. Bin width = 200 ms; see Fig. S1B for other bin widths. Insets show sample UMAP embeddings of population rate vectors for the first (top) and last (bottom) second of stimulation with 120 Hz vibration with different stimulus intensities (color); see Fig S2A for additional stimulus frequencies. UMAPs illustrate that rate vectors for different stimulus intensities are more linearly separable after adaptation, which supports better decoding. **C.** Same as B, but for decoding stimulus frequency. There was no sustained improvement in the discrimination of stimulus frequency from firing rate. See Fig. S2B for UMAPs for additional stimulus intensities.

Adaptation improved decodability of stimulus intensity from the population rate vector as evident from the sustained increase in decoder accuracy and mutual information (**Fig. 2B**). This was largely driven by improved discrimination of moderate intensity stimuli (**Fig. S1A**) and occurred for bin widths of 200-500 ms, but not for 100 ms-wide bins (**Fig. S1B**), most likely because narrow bins contain too few spikes (although this problem would be mitigated for larger populations of neurons). Improved decodability is visually evident in the UMAP embedding of population firing rates insofar as rate vectors for different stimulus intensities are more linearly separable in later intervals (**Figs. 2B insets** and **S2A**). Errors in decoding could be due to poor generalization (caused by differences between the training and test sets) or to poor coding (caused by overlapping rate distributions for different stimulus intensities) (54). To tease these apart, we repeated analysis using an SVC but now with a test set identical to the training set, and found the same sustained improvement in decodability of stimulus intensity (**Fig. S3A**), thus verifying that adaption improves coding. Comparable improvements were also evident using a random forest decoder (**Fig. S3B**) or linear decoder (**Fig. S3C**) instead of an SVC.

Unlike for stimulus intensity, rate coding of stimulus frequency improved modestly at first but eventually returned to baseline (**Fig. 2C**; see also **Figs. S1C,D** and **S2B**). Accuracy and mutual information remained mostly unchanged when tested on the same data set used for training (**Fig. S3D**) or when tested with a random forest decoder (**Fig. S3E**), and even dropped when tested with a linear decoder (**Fig. S3F**). These results suggest that rate coding of stimulus frequency does not consistently benefit from adaptation.

### The improvement in rate coding relies on nonuniform adaptation effects

We hypothesized that adaptation’s ability to improve rate-based coding of stimulus intensity emerges from population rate vectors for different stimulus intensities becoming more separable due to nonuniform (i.e. stimulus-dependent) effects of adaptation on LTMR tuning (Fig. 1C). To test this, an artificial dataset was created in which the tuning curve for each LTMR was linearly re-scaled (see Methods and **Fig. 3A**). In brief, for each neuron, entrainment was reduced from its starting value by the same amount across all stimuli; the reduction was calculated separately for each neuron and for each second-long interval based on the average change in entrainment for the interval of interest relative to the initial interval, in that neuron. Because each neuron’s rate is scaled separately, simulated population rate vectors are not a trivial re-scaling of vectors from the initial interval; instead, each dimension of the vector (i.e. each neuron) is scaled independently. The simulated linear adaptation yielded no improvement in rate-based coding of stimulus intensity (**Fig. 3B**) or frequency (**Fig. 3C**). In short, nonlinear adaptation effects are necessary for rate coding to benefit from adaptation.

**Figure 3.**
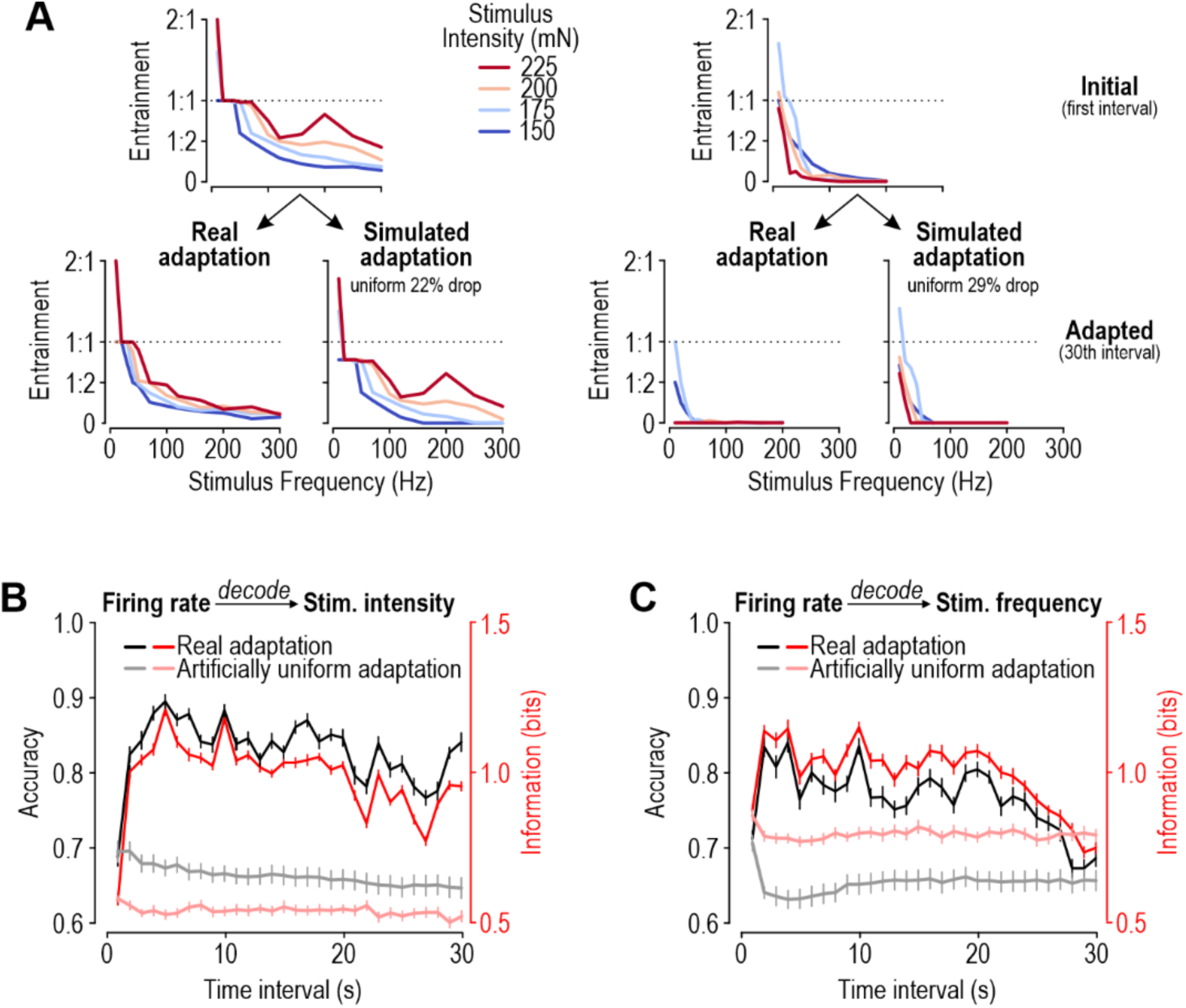
Improved rate coding of stimulus intensity relies on nonuniform adaptation effects. **A.** Sample tuning curves from two LTMRs initially (top) and after adaptation (bottom left). In the real dataset, the decrease in firing rate due to adaptation varies across stimulus parameters (see Fig. 1C). To simulate adaptation that is uniform across stimuli (bottom right), the initial tuning curve was linearly re-scaled by a percentage calculated by averaging the adaptation-mediated change in entrainment across all stimuli for the interval of interest relative to the first interval. **B.** Decodability of stimulus intensity from firing rate was not improved by artificially uniform adaptation (pale curves). Data for real adaptation (from Fig. 2B) are plotted in dark for comparison. Analysis is the same as for Fig. 2. **C.** Same as B but for decoding of stimulus frequency. Data for real adaptation (from to Fig. 2C) are replotted in dark for comparison.

### Adaptation improves the efficiency of encoding stimulus frequency with inter-spike intervals

Previous work suggests that LTMRs can encode vibration frequency using inter-spike intervals (ISIs) (8, 46, 47). As illustrated in **Figure 4A**, when driven by a single-harmonic periodic stimulus, LTMRs often spike at a consistent phase of the stimulus (see below) and those phase-locked spikes occur at integer multiples of the stimulus period; for example, a 200 Hz stimulus will drive spikes at integer multiples of the 5 ms stimulus period (5, 10, 15 ms, etc.). To represent the ISIs of an LTMR population, we calculated ISIs from each neuron’s spike train during a 100 ms bin, and then pooled these ISIs across the population to create a distribution (**Fig. 4B**). We then trained an SVC decoder to predict either stimulus frequency or intensity from this ISI distribution.

**Figure 4.**
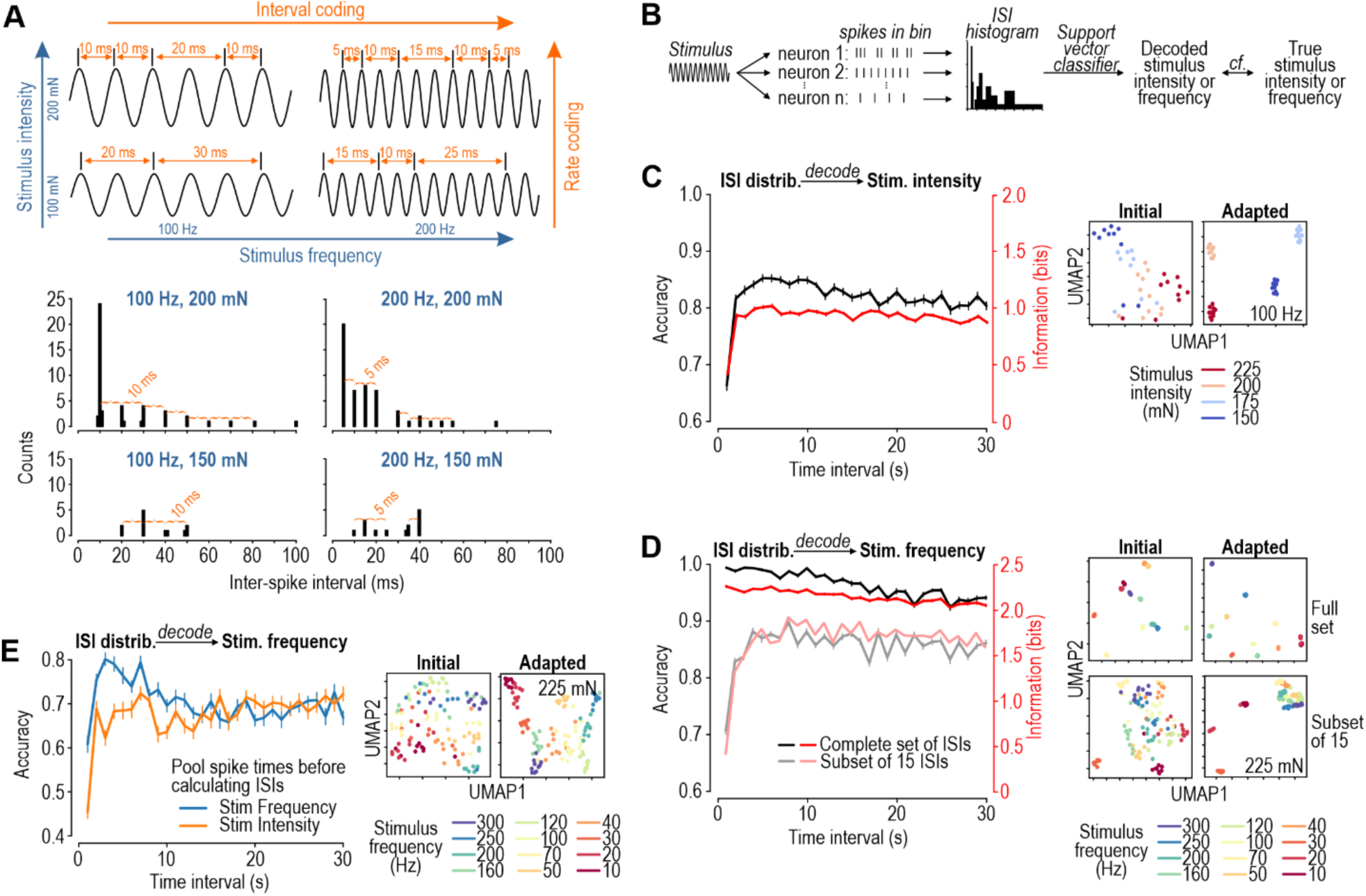
Adaptation improves the efficiency of temporal coding of stimulus frequency. **A.** Stimulus intensity and frequency can be simultaneously encoded using the rate and timing (i.e. interval between) spikes, respectively. ISIs occur at integer multiples of the stimulus period by occurring at a consistent phase of periodic stimuli but not necessarily on every cycle; entrainment varies with stimulus intensity. Histograms (bottom) show sample ISI distributions for different stimuli. Distributions were generated from real data by randomly choosing spike times in a 100 ms interval from the 30th second of each neuron’s response and then calculating ISI values independently for each of 26 neurons before pooling. **B.** Procedure for evaluating population temporal coding. A different ISI distribution was created for each 100 ms interval of each stimulus presentation by calculating ISIs in each neuron, and then pooling across the population (see Methods). The 21 edges of the histogram were logarithmically spaced to better represent smaller ISIs while keeping dimensionality small. Stimulus parameters were then decoded from this 20-dimensional vector using an SVC. **C.** Decoding of stimulus intensity from ISI distributions, using all ISIs. Accuracy (black) and mutual information (red) both exhibited a sustained increase. A different classifier was used for each second of stimulation. The train/test split (0.7/0.3) was repeated 20 times with different splits. UMAPs illustrate that ISI distributions for different stimulus intensities are more linearly separable after adaptation. **D.** Same as C, but for decoding stimulus frequency from ISI distributions using all ISIs (dark curves) or subsets of 15 ISIs (pale curves). When using all ISIs, accuracy and mutual information both start high and decrease slightly over time. In contrast, when using an equivalent number of ISIs from all intervals by subsampling ISIs, a sustained increase in performance was observed. See Fig. S4B for effects of subsampling different numbers of ISIs. These results show that encoding is improved by using more spikes, but adaptation increases the information conveyed by each spike; however, the latter effect can go unnoticed if encoding is already near-perfect (i.e. there is a ceiling effect). Comparing UMAPs when using all ISIs (left) or when subsampling a fixed number of ISIs (right) shows that adaptation improves discrimination in the latter case (when initial encoding is poorer). **E.** Same as dark data in D, but using (all) ISIs calculated from spike times pooled across all neurons (instead of calculating ISIs from spike times in each neuron, and then pooling ISIs). There is a sustained improvement in the decoding of both stimulus intensity and frequency. Notably, accuracy of stimulus frequency decoding is lower than in panel D because calculating ISIs from the pooled spikes yields fewer informative ISIs, which in turn prevents a ceiling effect. UMAPs illustrate the increased discriminability of stimulus frequency after adaptation.

Decoding of stimulus intensity from the ISI distribution improved rapidly and remained above baseline during the 30-s-long response (**Fig. 4C**). Similar to the rate code, this improvement was mostly driven by better decoding of moderate intensity stimuli (**Fig. S4A**). Since the reciprocal of the average ISI corresponds to firing rate, the improvement seen here is expected from the improvement already reported in Fig. 2B. However, unlike with the rate code, the ISI distribution enabled near-perfect decoding of stimulus frequency (**Fig. 4D**), consistent with an ISI-based temporal code being better able to represent stimulus frequency. Notably, encoding of stimulus frequency did not, on first inspection, benefit from adaptation insofar as decodability decreased slightly over the 30-s-long response (**Fig. 4D, dark curves**) but, notably, there is little room for adaptation to convey any benefit given how good decoding is to begin with.

Recent work has suggested that metabolic efficiency, or bits/spike, is a more potent constraint on neural design than coding efficiency, or bits/second (55, 56). Notably, the number of spikes is high initially, which could support good but metabolically inefficient coding. To investigate this, we uniformly subsampled an equal number of ISIs for each interval before decoding (see Methods). Decoding of stimulus frequency from distributions of 15 ISIs improved substantially after the first second of the response (**Fig. 4D, pale curves**). Similar improvement was evident for decoding from distributions comprising relatively few (≤15) ISIs (**Fig. S4**); decoding from larger subsampled ISI distributions revealed the same ceiling effect seen when using all ISIs. Subsampling spike-times before calculating ISIs (as opposed to subsampling ISIs after calculating them from the original spike trains) yielded similar results (**Fig. S4D,E**). Furthermore, rather than calculating ISIs for each neuron before pooling them, we also investigated pooling spike times across the population and then calculating ISIs (which more accurately represents input to a downstream neuron receiving convergent input from multiple co-activated LTMRs). With the latter representation, there was a sustained improvement in the decodability of stimulus intensity and frequency (**Fig. 4E**). The benefit of adaptation on decoding of stimulus frequency was evident despite using all ISIs because pooling spike times before calculating ISIs yielded poorer performance overall, thus preventing a ceiling effect. Results were consistent across different decoders (**Fig. S5**).

Overall, these results confirm that an ISI-based code is better suited for representing stimulus frequency than a rate code, and also demonstrate that the metabolic efficiency of this code improves with adaptation.

### The improvement in temporal coding efficiency relies on improved phase-locking precision

Interval-based coding relies on precisely phase-locked spikes lest the ISI pattern become indiscernible (**Fig. 5A**). Phase-locking can be represented by plotting spike times relative to the phase of the periodic stimulus (see polar plots on **Fig. 5A**). Variance of that polar distribution is inversely related to phase-locking precision (see Methods). Phase-locking can be reduced by jittered spike times (evident as a broadened distribution) or because >1 spike occurs per stimulus cycle (causing a multimodal distribution), where “additional” spikes tend to delay the first spike on the net cycle, and so on (see cartoon on top panel of **Fig. 5A**). Regardless of the mechanism, phase-locking tended to improve with adaptation (**Fig. 5B**). Notably, “additional” spikes were excluded from jitter calculations, but their inclusion would only exaggerate effects described below.

**Figure 5.**
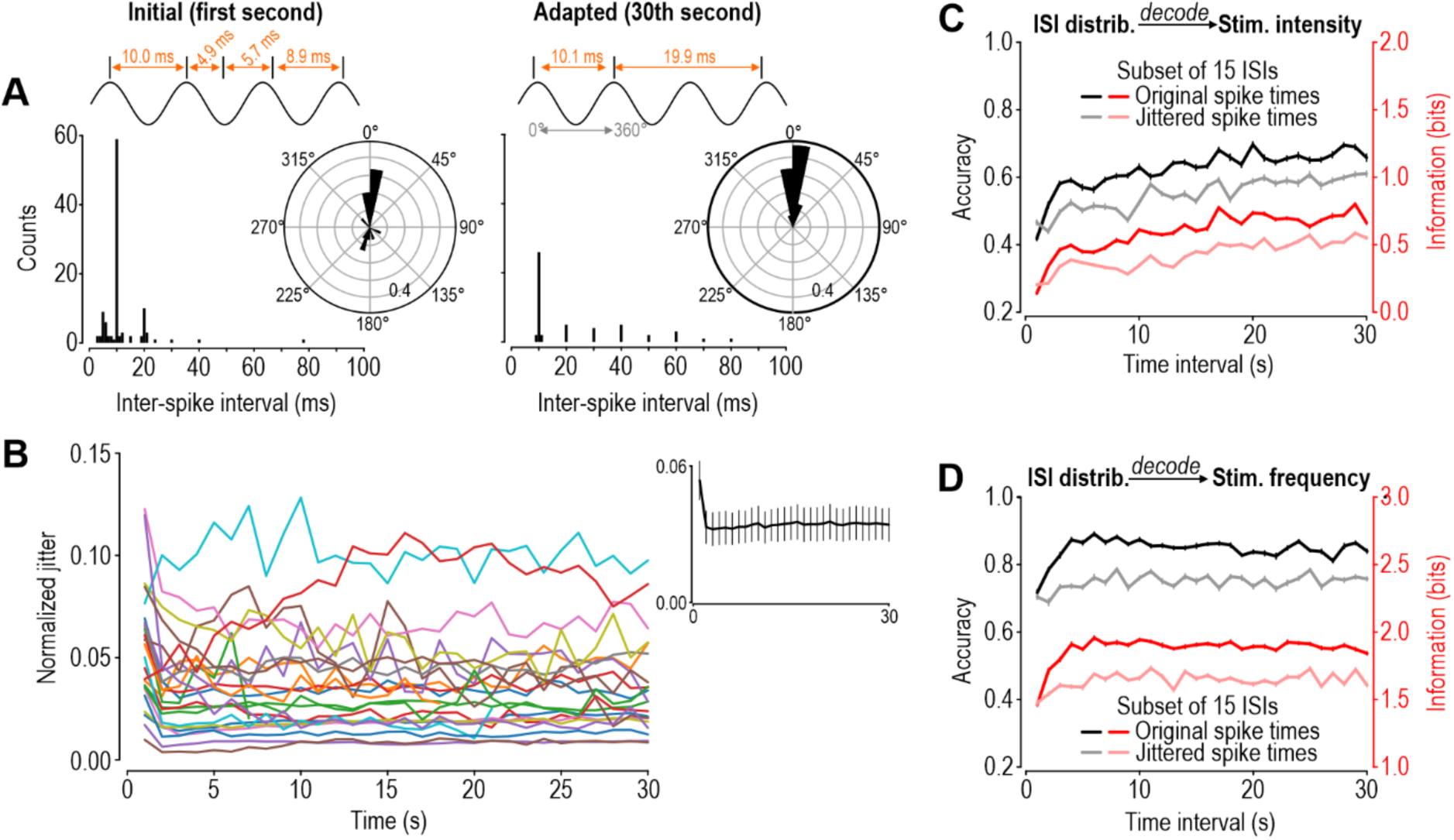
Precisely phase-locked spikes are required for temporal coding of stimulus frequency. **A.** Sample ISIs distribution from response to 100 Hz, 200 mN vibration. Cartoons at top show spike times relative to the periodic stimulus and the resulting ISIs. After adaptation (30^th^ interval, right), all spikes occur at nearly the same stimulus phase (see polar plot), resulting in ISIs clearly distributed at integer multiples of the stimulus period. Before adaptation (first interval, left), phase-locking is poorer, in part because >1 spike occur on some stimulus cycles; the resulting ISI distribution is less obviously derived from 100 Hz stimulation. These ISI distributions were created by calculating ISIs for each neuron, and then pooling ISIs across the population, as opposed to pooling spike times and then calculating ISIs. Polar plots were generated by subsampling 15 ISIs from each distribution and converting them to angular coordinates (see Methods). **B.** Normalized variance in polar plots for each of 26 neurons (different colors). Polar plots were analyzed for each stimulus separately, and then their normalized angular variance (divided by 2π) was averaged across stimuli. This was done separately for each second-long interval of the response. Inset shows average jitter (± SEM) across all neurons. **C.** Impact on decoding stimulus intensity of jittering times of all spikes occurring after the first second by 6% of the corresponding stimulus period (pale curves) for comparison with original data (dark, from Fig. 4C). **D.** Same as C, but for decoding of stimulus frequency.

We hypothesized that the increased efficiency of interval-based coding of stimulus frequency (see Fig. 4) emerges from the adaptation-mediated improvement in phase-locking precision. Notably, without subsampling, the greater number of ISIs during the initial response might offset the reduced precision of individual spikes, consistent with the ceiling effect shown in Fig. S4. To directly test if phase-locking precision is important for temporal coding of stimulus frequency, we artificially manipulated spike times in all but the first 1-s interval to nullify the improvement reported in Figure 5B; specifically, we artificially jittered spike times by 6% of the stimulus period (see **Fig. S6** other degrees of jittering). Jittering modestly impaired encoding of stimulus intensity without qualitatively altering the change in encoding over time (**Fig. 5C**) but, as predicted, it completely abolished the improvement in encoding of stimulus frequency over time (**Fig. 5D**).

Overall, our results suggest that improvements in phase-locking precision mediated by adaptation allow each spike (or the ISI it forms) to convey more information, thus increasing metabolic efficiency of the temporal code. This effect is not evident when there is an abundance of ISIs (which is usually true for the initial response) because excellent coding is achieved with many poorly timed spikes. The adapted response achieves excellent coding with fewer, well-timed spikes.

### Adaptation improves decodability of stimulus identity (both intensity and frequency)

Ultimately, spike trains encode stimulus intensity and frequency concurrently. Testing decodability of both stimulus features confirmed that adaptation increases the decodability of both features from the population firing rate (**Fig. 6A**) and from the ISI distribution (**Fig. 6B**). Note that the ISI-based temporal code contains more information overall than the rate-based code. To summarize, despite adaptation causing reduced spiking, the remaining spikes form better (i.e. more decodable) representations of both stimulus intensity and frequency.

**Figure 6.**
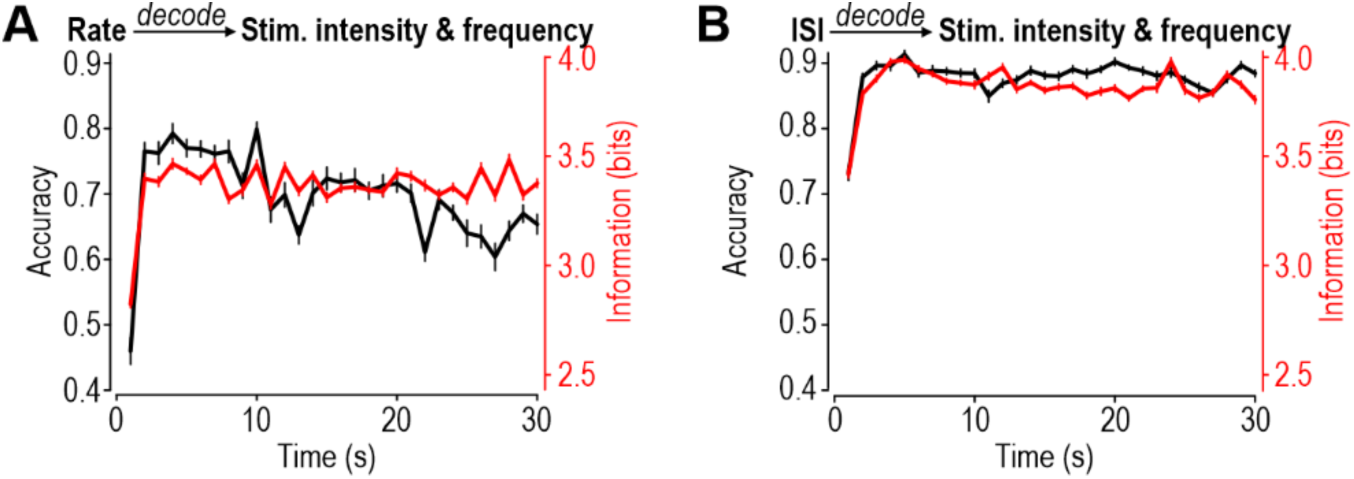
Adaptation improves encoding of stimulus identity (i.e. intensity and frequency). **A.** Decoding both stimulus intensity and frequency simultaneously from the population firing rate using same methodology as in Figure 2, with 200 ms bins. **B.** Decoding of stimulus intensity and frequency from ISI distributions using the same methodology as in Figure 4, namely using all ISIs without subsampling or jittering. Temporal coding is significantly more accurate and informative than rate coding.

## DISCUSSION

The first description of firing rate adaptation was reported from recordings of primary cutaneous afferents almost a century ago (57) but the implications for somatosensory coding have gone unexplored. Our results show that adaptation in LTMRs improves the neural representation of somatosensory input to the CNS (**Fig. 6**). This was true for both rate and temporal coding. Rate-based coding of stimulus intensity especially benefited from adaptation (**Fig. 2**). This improvement relied on nonuniform changes in firing rate, meaning responses to different stimulus parameters change differently, which resulted in tuning curves changing shape as opposed to being re-scaled linearly (**Fig. 3**). Such changes translate to population rate vectors for different stimulus intensities becoming more discriminable, as visually evident from UMAP embeddings. ISI-based coding of stimulus frequency was extremely good (**Fig. 4**), so much so that adaptation-mediated improvements were occluded by a ceiling effect. However, deeper analysis revealed that adaptation increases the information conveyed by each ISI, meaning adapted responses achieve good coding with fewer, well-times spikes, unlike initial responses that achieve good coding with many poorly timed spikes. This improvement in temporal coding derives from spike timing (i.e. phase-locking) becoming more precise with adaptation (**Fig. 5**). Overall, our results show that adaptation can improve somatosensory encoding, but different coding strategies benefit via different mechanisms.

Before the onset of stimulation, a neuron is primed for stimulus detection (using high gain). But if stimulation is maintained, the neuron adjusts its gain for the benefit of stimulus discrimination. Previous reports have suggested adaptation trades off absolute stimulus information for relative discriminability of similar stimuli (35, 58). Information about absolute stimulus intensity is not necessarily lost and may instead be tracked by slower statistical features of the neural response (22). With our testing protocol, we observed mostly decreases in spike count over time, but spike counts can increase with adaptation under other conditions (22, 25, 59, 60).

Mechanosensory adaptation has explicit consequences for perception in humans. Most relevantly, Tommerdahl et al. found that adaptation to a high-frequency stimulus (200 Hz) improved subsequent performance in frequency-discrimination tasks for high but not low frequencies, while adaptation to a low frequency stimulus (20 Hz) improved discrimination of low but not high frequencies (61). Our work suggests that the neural correlate of behavioural adaptation might start much earlier in the mechanosensory pathway than S1 cortex, where adaptation has already been normatively investigated (29, 34). Similarly, in the visual system, adaptation has been observed not just in V1 (23, 33–35), but also in retina (27, 28). Adaptation in early visual neurons has perceptual consequences (24, 56). It is thus plausible that adaptation in primary mechanosensory afferents also impacts perception. Primary sensory neurons cannot be thought of as purely passive relays but, instead, operate as dynamically self-optimizing feature extractors (25, 26, 62).

The mechanism of adaptation is likely context-dependent. While there has been evidence for circuit-level adaptation (25, 26, 63), the lack of synaptic connectivity among LTMRs means adaptation at this stage arises from changes intrinsic to each neuron. At a computational level, single-cell adaptation has been described as an active inference process, where a cell leverages its metabolic computational capacity to track changes in the stimulus distribution, and adjust its encoding function accordingly (60, 64, 65). In support of this, a cell adapts faster when a change in the stimulus distribution is more discriminable (60), and cells integrate evidence at multiple timescales simultaneously (22, 34, 59, 60, 66, 67). Like the multiplexing of codes, multiple timescales of adaptation can introduce additional information channels which encode different dynamical timescales of the stimulus (22, 25, 60, 64). While our data only reports one timescale of adaptation, future work should use dynamically richer stimuli to probe for multiple timescales of LTMR adaptation.

While normative descriptions of adaptation are useful, a descriptive modelling approach is necessary to understand its algorithmic implementation (2). Fits of general linear-nonlinear models have reported changes to both the stimulus filter and the nonlinearity with adaptation (23, 26, 64, 68). Recent work has suggested that many features of adaptation can be described through a simple process, where neurons take the fractional derivative of their stimuli (59, 69). This allows neurons to be both sensitive to fast changes, (encoded with fast spiking dynamics) while also integrating slow changes (encoded with slow spike-train dynamics). However, it is still debated whether such models are expressive enough to capture the computational complexity of the adaptive inference process (60). For example, the sensitivity of adaptation to higher-order moments of a stimulus distribution has not been captured by algorithmic models, such as fractional differentiation, but has been experimentally observed (60). The appropriate modelling of a neuron’s dynamical input-output function with respect to its receptive fields, across multiple contexts, might require more expressive approaches such as artificial neural networks (ANN) (65, 70–72). While ANN-based models would be less interpretable and possibly have less research utility (26), the increased predictive power they provide would be more suitable for brain-computer interfaces such as next-generation prosthetics.

At the level of biological implementation, various adaptive mechanisms have been explored, such as the interaction of multiple after-hyperpolarization currents (25, 73–77). To account for the diversity of adaptation timescales in a single neuron, these multiple currents should be rapidly tunable by the inference program (60). Other mechanisms are conceivable. The adaptation observed in LTMRs does not involve synaptic mechanisms (see above) but might involve changes in mechanotransduction through adaptation in mechanosensitive channels (e.g. piezo) or how skin transmits force to those channels.

In conclusion, we report that LTMRs adapt to vibrotactile stimulation in such a way as to improve the discriminability of similar stimuli. This occurs due to changes in the rate and temporal encoding of stimulus intensity and frequency, respectively. Notably, while adaptation in LTMRs increases information conveyed to central neurons, our approach does not demonstrate that central neurons capitalize on that improvement. We demonstrated that different types of decoders can reveal the benefits of adaptation on encoding, which increases the likelihood that a biological decoder could likewise benefit, but we did not test specific biological decoders. Psychophysical tests are important in that regard, but it is difficult to ascribe improved psychophysical discrimination to adaptation at specific points in the neuraxis. Our results suggest that adaptation in LTMRs contribute to enhanced discrimination. Future work should study LTMRs using a wide range of intelligently sampled, dynamic, and repeated stimuli (78), ranging from white noise to naturalistic multi-harmonic textures (79). Deeper understanding of natural neural coding strategies and how they benefit from adaptation may, ultimately, have practical implications for optimizing next-generation prosthetic limbs that incorporate sensory feedback (45, 80).

## METHODS

### In vivo electrophysiology

All procedures were approved by the Animal Care Committee of the Hospital for Sick Children (protocol #47072). Adult (200-350 g) male Sprague-Dawley rats were obtained from Charles River, Montreal. Detailed methods are described by Medlock et al. (81). In brief, rats were anesthetized with 1.2 g/kg urethane i.p. and received 10% top-ups as needed. The L4 and L5 dorsal root ganglia (DRG) were exposed by laminectomies and, with the animal stabilized in a stereotaxic frame (Narishige), a multielectrode array (NeuroNexus, A4 type) was inserted in either DRG. Signals were amplified, digitized at 40 kHz and high-pass filtered at 300 Hz using an OmniPlex Data Acquisition System (Plexon). Precisely controlled vibrotactile stimulation was applied to the ipsilateral hind paw using a model 300C-I mechanical stimulator (Aurora Scientific) with the flat-tipped indenter (1 mm diameter) positioned on the paw with a micromanipulator. Thirty-second long sinusoidal vibrations of different intensities (50-225 mN) and frequencies (10-300 Hz) were applied while recording each unit. Stimuli were controlled by a Power1401 A-D board and Signal v5 software (Cambridge Electronic Design). Stimulus timing was aligned to recorded responses using triggers sent from Signal to Omniplex. Single units were isolated using Offline Sorter v5 Software (Plexon) and analyzed with MATLAB (MathWorks, R2022) and Python 3.10.

### Tuning curves

The spike train recorded from each neuron during each vibrotactile stimulation was divided into thirty 1-sec-long intervals. A different tuning curve was created for each interval, although we only show tuning curves for the first and final (30^th^) interval. Tuning curves report %entrainment for each frequency-intensity combination, revealing a neuron’s rate-based encoding function. This was calculated as:

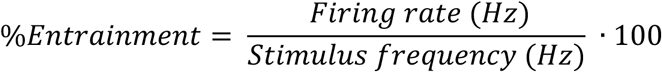

### Measuring a lower-bound on mutual information from population rate codes

Population rate codes were represented as vectors constructed from the firing rate of each neuron in the population, meaning vector length corresponds to the number of constituent neurons. The spikes produced by each neuron must be counted over some time window (bin); we tested bins of length 100, 200, 333 and 500 ms (Figure S1). We did not use a sliding window, and so a 200 ms window means 1 second of data was divided into five consecutive bins. By not using a sliding window, we avoided overlap that would produce highly similar rate vectors, which might in-turn optimize for trivial decoding strategies (see below).

A support-vector classifier (SVC) was used to decode either the stimulus intensity or frequency from the population rate vector at any given time. All classification was done using the scikit-learn package in Python, with default parameters except C=4 for decoding intensity and C=10 for decoding frequency. We also tested a linear decoder, using the scikit-learn LogisticRegression with max_iter set to 1000 and penalty (regularizer) set to None, and a random forest classifier from the same Python package, using its default parameters.

The SVC was applied to a dataset comprising 8 neurons tested with 10 frequencies (10-200 Hz) repeated at 4 intensities (150-225 mN). This was the largest subset of the full dataset without any missing trials. Missing trials were due to either loss of a neuron during a long recording session or unreliable spike sorting after-the-fact. The inclusion of neurons with missing trials might optimize for trivial decoding strategies. The size of the dataset depends on the integration window used; splitting a one second interval of data into 200 ms bins yields 200 different rate vectors (number of frequencies [10] x number of intensities [4] x number of 200 ms bins in a second [5]). When using 100 ms bins, there were 400 vectors in the dataset, etc.

To measure how well rate vectors encode stimulus intensity, we split data based on the second-long interval in which it was recorded (1-30 inclusive) and then applied 200 ms-long bins (other bin lengths were tested in Fig. S1). A different classifier was trained for each interval. Since there are four different intensities, there are 50 training samples per intensity. We used 70% of each dataset to train a classifier to predict stimulus intensity from these vectors, while the remaining 30% test set was used to calculate accuracy and a lower-bound on mutual information (see below). This was repeated 20 times, each time with a different train/test split, allowing for a mean and standard-error-of-the-mean (SEM) to be calculated. The same was done when decoding stimulus frequency, except in that case there are only 20 training examples per class, since there are 10 frequencies to be predicted.

Since classifiers can output probability distributions over their output targets for each test example, a lower bound for mutual information between stimulus identity and neural response can be calculated. This was done by subtracting the entropy of this output distribution from the entropy of the prior (2 bits for intensity [=log2(4) based on 4 uniformly tested intensities] and 3.32 bits for frequency [=log_2_(10) based on 10 uniformly tested frequencies]). Averaging this difference in entropy over every sample in the test-set would result in the mutual information for one train/test split. To validate mutual information measures, decoding accuracy was also measured for 20 different train/test splits to yield mean ± SEM values. Accuracy was calculated as the number of correct predictions divided by the total number of predictions.

### UMAP plots

UMAP plots were created using the Python package umap-learn. For every plot made throughout the paper, the default parameters (15 neighbours and a minimum distance of 0.1) were used.

### Generating simulated firing rate data

To test the hypothesis that adaptation improves rate coding by affecting firing rate differently for each stimulus (i.e. non-uniformly), we generated an artificial dataset in which adaptation was stimulus-independent for each neuron. This might reasonably occur if adaptation was purely a fatigue-based process. To generate the dataset, each second-long interval of the 30 s response was compared to the first interval, and the average change in stimulus entrainment was calculated over stimuli in each neuron. For example, in neuron 50, entrainment fell by an average of 22% in the last second relative to the first second, so to generate the simulated tuning curve for the last second, 22% was subtracted from the entrainment values of the tuning curve for the first second. Entrainment values were rectified to avoid negatives. Firing rates used to make population rate vectors were then scaled by the new simulated entrainment value for each stimulus frequency. This scaling was via element-wise multiplication, as each neuron had a different decrease in entrainment for that stimulus and second of stimulation. Simulated firing rate data were then put into the same pipeline used to analyze real data (see above).

### Measuring Information available from interspike intervals (ISIs)

Phase-locked spikes occur at ISIs that correspond to integer multiples of the stimulus period, thus producing spiking patterns that depend on (i.e. encode) stimulus frequency. Beyond the temporal encoding of stimulus frequency, the *average* ISI depends on stimulus intensity because stimulus intensity modulates entrainment. In other words, stimulus frequency dictates the spacing between individual peaks on the ISI histogram, but stimulus intensity dictates the central tendency of the overall distribution. The decoder described below capitalizes on ISI patterning and average ISI to extract a lower bound on mutual information.

Like for rate coding (see above), the quality of temporal coding was assessed based on ability to infer stimulus parameters from the response. Ideally, a population temporal code would be represented by a high-dimensional binary matrix, where rows represent different neurons and columns represent short (1-2 ms) intervals in which a single spike may or may not occur, but we did not have sufficient data to train decoders from such representations. Instead, to represent information at 1-2 ms resolution while keeping the dimensionality manageable, we used a 20-bin representation of the ISI distribution. When constructing the ISI distribution, histogram edges were logarithmically spaced to better represent small intervals (21 edges: 1.0, 1.3, 1.6, 2.0, 2.5, 3.2, 4.0, 5.0, 6.3, 7.9, 10.0, 12.6, 15.8, 20.0, 25.1, 31.6, 39.8, 50.1, 63.1, 79.4, 100 ms). This 20-element vector was then input to an SVC to decode either stimulus frequency or intensity. For most figures, ISIs were calculated for each neuron’s spike train, and then pooled across the population; in other cases, spike times were pooled across the population, and then ISIs were calculated. The final distribution was always normalized to sum to 1.

The ISI distribution was constructed from the ISIs of all 26 neurons during the same 100 ms period, and this was done across 12 stimulus frequencies (10-300 Hz) and 4 intensities (150-225 mN). This is unlike the rate code analysis, which used only 8 neurons and 10 frequencies to avoid the inclusion of neurons with missing trials. While missing trials would directly affect the population rate vectors in a stimulus-specific way, they are less likely to have a strong effect on the *pattern* of the ISI distribution.

When splitting one second of data into 100 ms bins, there were 480 different population ISI vectors (12×4×10) per second. The train/test split was repeated 20 times. The lower bound on mutual information was calculated using the same strategy, except now the maximum entropy of the frequency is 3.58 [=log2(12)], accounting for the additional two frequencies being predicted (250 and 300 Hz). The SVC classifier had a radial basis kernel and C=10 for both intensity and frequency.

### Measuring phase-locking precision

ISI patterning requires spikes to occur at a consistent phase of the period stimulus (i.e. phase lock). To measure this, we computed the variance in phase-locking precision. The spike times were first converted from units of time to units of radians, depending on the phase of the stimulus. Only the first spike was considered in instances where a neuron fired >1 spike per stimulus cycle. The converted spike times were used to create a polar distribution, with normalized variance of the distribution being calculated as:

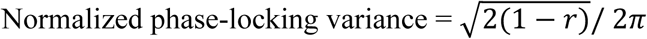

Where *r* is the resultant vector of the polar distribution. This value is inversely proportional to the degree of phase-locking precision.

### Subsampling ISIs and subsampling spikes

We subsampled, without replacement, a maximum number of ISIs using two strategies. In the first strategy, we created the pooled ISIs and then sub-sampled up to the designated number of ISIs; if the designated number exceeded the number of available ISIs, nothing changed. In the second strategy, spike times were subsampled from the population until the designated number of ISIs was reached, and ISIs were calculated from the timing of pooled spikes. To do this, we created a binary matrix with each row representing a neuron, and each column representing the presence or absence of a spike in a 100 ms interval discretized at 1 kHz. The total number of ISIs that could be calculated from this population matrix can be computed; each neuron will have n-1 ISIs, where n is the number of spikes the neuron fires. Therefore, total ISIs can be calculated by summing across the columns, subtracting 1 from each element, and then summing across the rows. A while loop was used to randomly remove spikes until there was a maximum number of ISIs from the spike matrix, and then those ISIs were calculated and used.

### Artificially jittering spike-times

To test if phase-locking is important for encoding stimulus frequency, we artificially manipulated the phase-locking precision of different trials. Since the average variance during the first second-long interval of stimulation was 0.06 (i.e. 6% of the stimulus period), we artificially jittered each spike time collected during subsequent intervals (2-30 s) by a standard deviation equal to 6% of the stimulus period. For each spike time, a random number was sampled from a Gaussian distribution. This random number was then multiplied by 6% of the stimulus period, and added to the spike time. We also tested other levels of jitter (see Fig. S6). This manipulation was done separately for each stimulus, based on its frequency and thus period. Informativeness of the artificially jittered spike train was calculated as described above.

### Measuring decodability of both stimulus intensity and frequency

We also used the SVC decoder to classify stimulus identity (10 frequencies repeated at 4 intensities), for a total of 40 classes to be predicted. When using the rate code, each second-long interval was divided into 200 ms bins, yielding a dataset of 200 vectors, or 5 per data class. When using the temporal code, each second was split into 100 ms intervals, yielding 10 vectors per class. The train/test split was repeated 20 times. The lower bound on mutual information was calculated using the same strategy, except now the maximum entropy of the frequency is 5.32 [=log2(40)]. The SVC classifier had a radial basis kernel and C=10 for both intensity and frequency.

## Acknowledgments

This work was funded by the Canadian Institutes of Health Research (CIHR) Foundation Grant 167276 to SAP.

## SUPPLEMENTAL FIGURES

**Figure S1.**
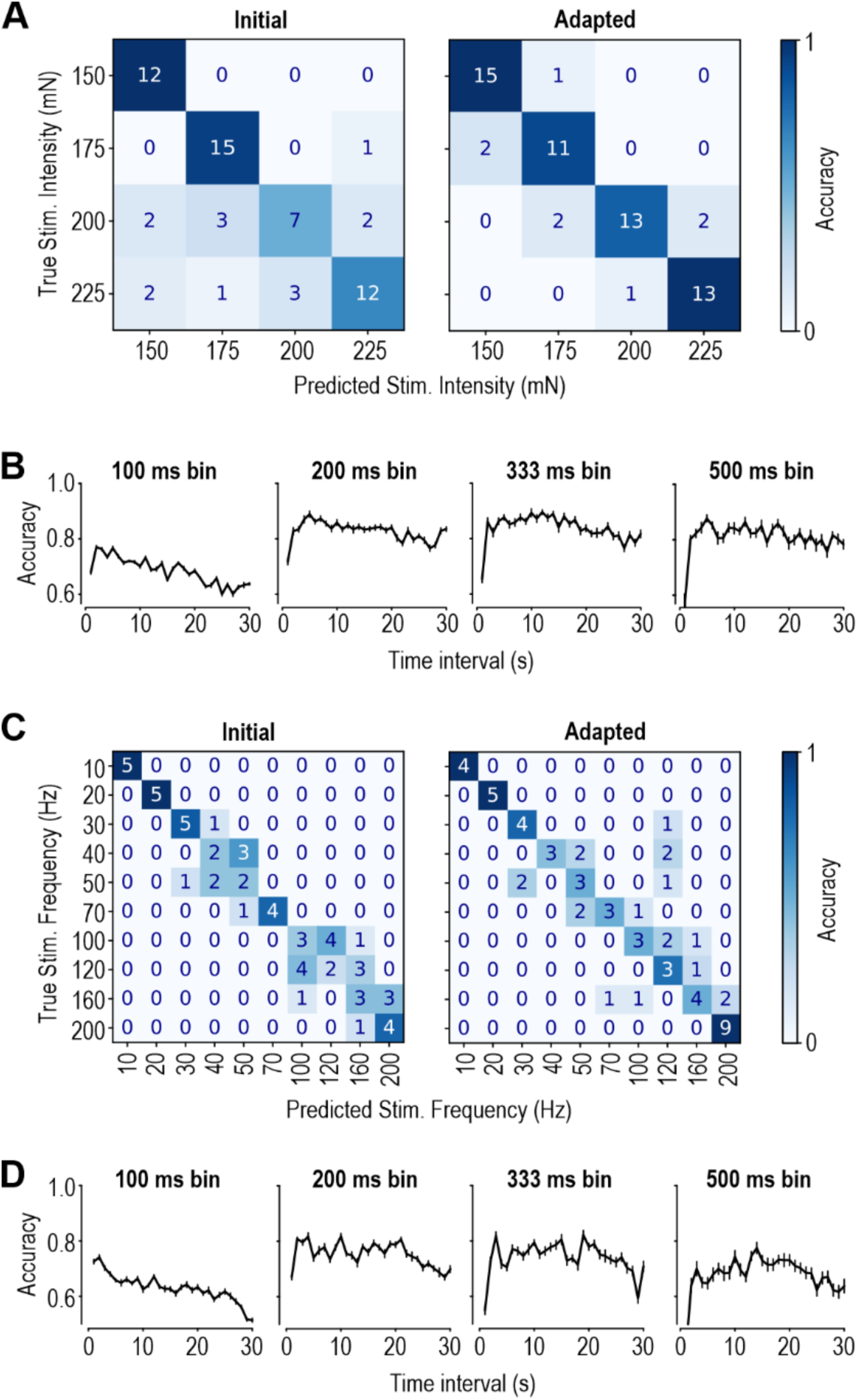
Extended analysis of decoding from population rate vectors. **A.** Correlation matrices for actual and decoded stimulus intensity based on data from the first second of stimulation (initial, left) compared to the 30th second (adapted, right). This is one of the 20 test sets generated for Fig. 2B. **B.** Accuracy of intensity decoding when spikes are counted in different width bins. The 200 ms integration window is the same data as in Fig. 2B. **C,D.** Same as A and B, but for decoding stimulus frequency. Compare to Fig. 2C.

**Figure S2.**
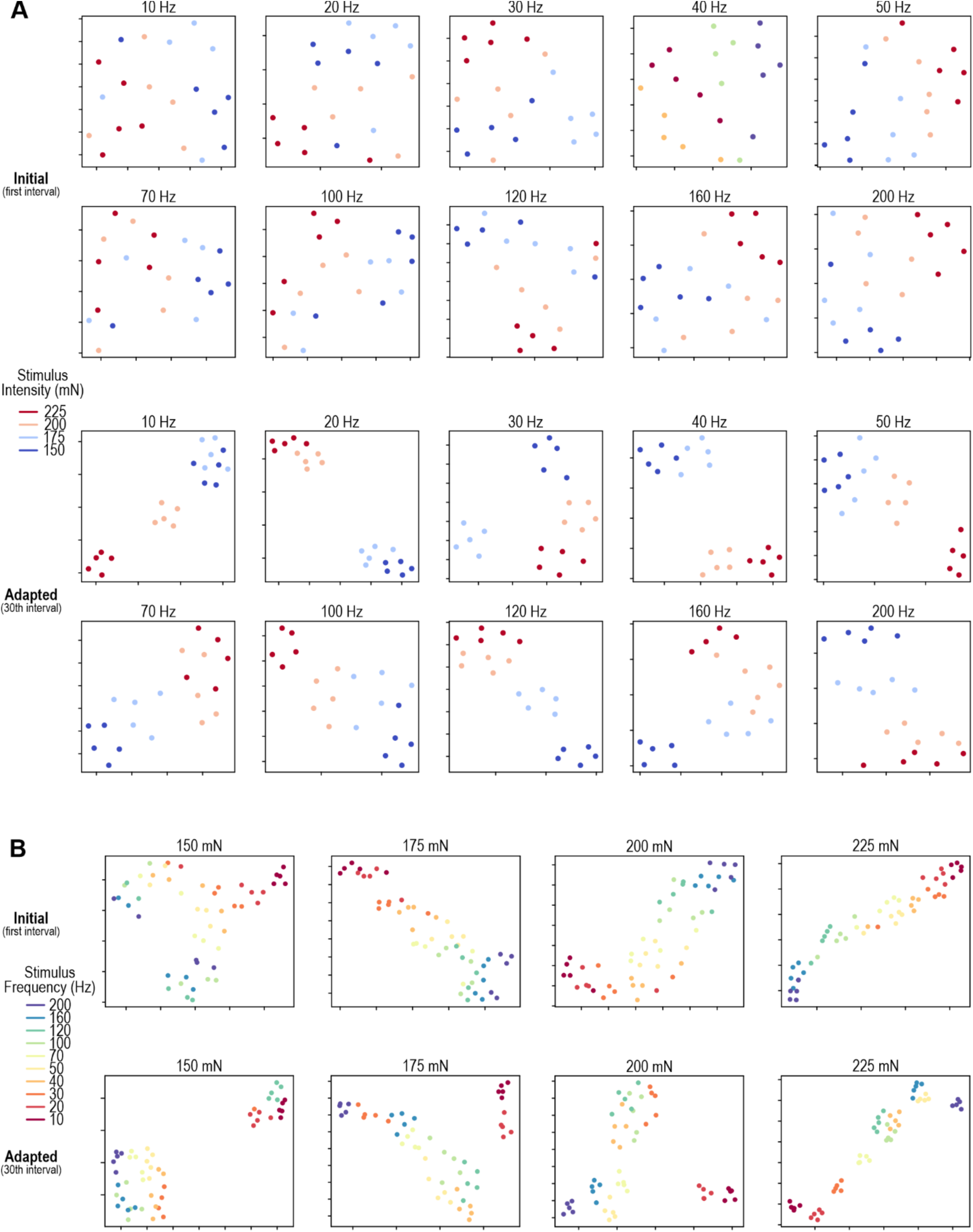
UMAP embeddings of population firing rate vectors before and after adaptation. **A.** UMAP embeddings of rate vectors for different stimulus intensities (color). Each panel shows rate vectors for a different stimulus frequency. Rate vectors for different stimulus intensities are more linearly separable after adaptation (bottom two rows) than initially (top two rows). **B.** Same as A but for rate vectors for different stimulus frequencies (color). Each panel shows rate vectors for a different stimulus intensity. Unlike in A, rate vectors for different stimulus frequencies are not more linearly separable after adaptation (bottom row) than initially (top row).

**Figure S3.**
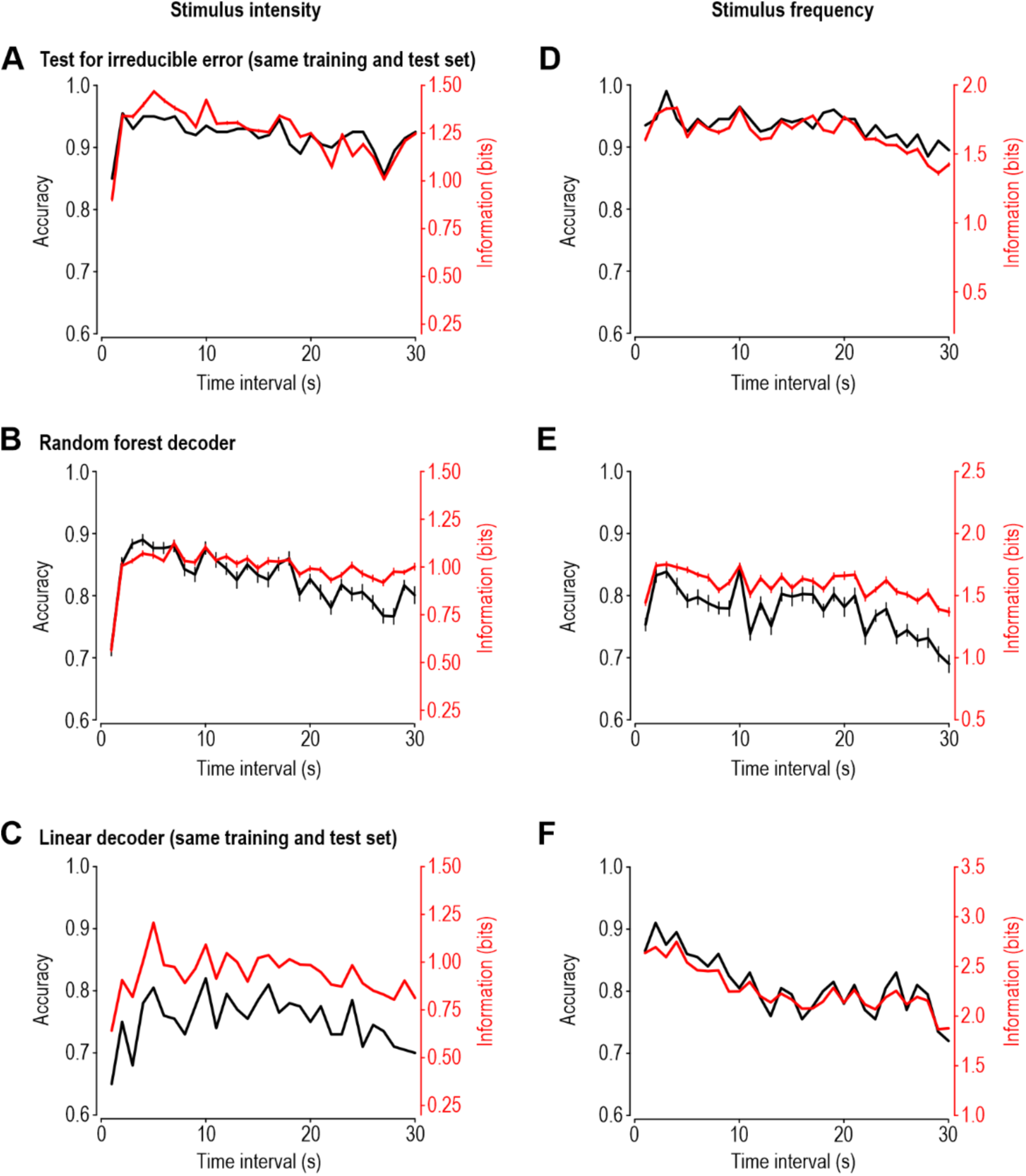
Comparison of decoders to assess discriminability of stimulus intensity (A-C) or frequency (D-F) from population firing rate. **A.** Same as Fig. 2B (decoding stimulus intensity) using an SVC but with the training set equal to the test set. This approach removes error due to failure of the model to generalize from the training set to the test set, and thus isolates the contribution of irreducible error (i.e. inability to discriminate different stimuli because they evoke similar firing rates). **B.** Same as Fig. 2B but using a random forest decoder instead of an SVC. The train/test split (0.7/0.3) is repeated 20 times with different splits. **C.** Same as Fig. 2B but using a linear decoder instead of an SVC. Training set is equal to test set. **D-F**. Same as Fig. 2C (decoding stimulus frequency) using an SVC but with the same training and test set (**D**), a random forest decoder (**E**), or a linear decoder (**F**). Regardless of the decoder, rate-based coding of stimulus intensity benefited from adaptation whereas rate-based coding of stimulus frequency did not.

**Figure S4.**
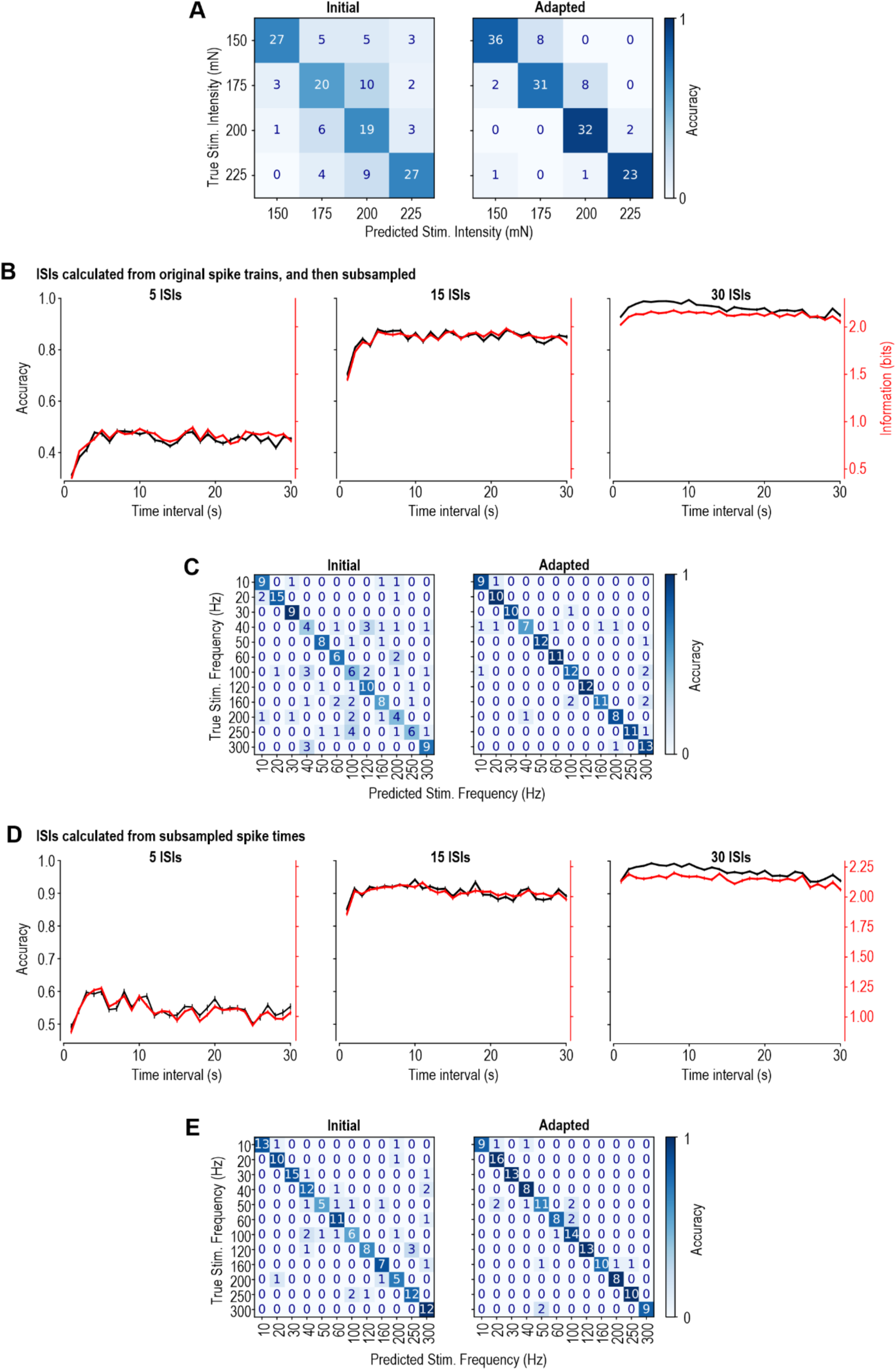
Extended analysis of decoding from ISI distributions. **A.** Correlation matrices for actual and decoded stimulus intensity based on data from the first second of stimulation (left) compared to the 30th second (right) for one of the 20 test sets used to generate Fig. 4C. **B.** Accuracy of frequency decoding when using different numbers of subsampled ISIs (5, 15, or 30 ISIs from left to right). Pale data in Fig. 4D corresponds to data for 15 ISIs shown here. **C.** Correlation matrices for actual and decoded stimulus frequency based on data from the first second of stimulation (left) compared to the 30th second (right) from one of the test sets using 15 subsampled ISIs, like in panel B **D)** Same as B, but instead of subsampling ISIs after calculating them, spikes times were subsampled to yield the target number of ISIs (5, 15, or 30), and ISIs were then calculated from subsampled spike times (see Methods). The same improvement in decoding is seen. **E)** Same as part C, but with the data from the middle panel of part D, when the subsample number is equal to 15 ISIs.

**Figure S5.**
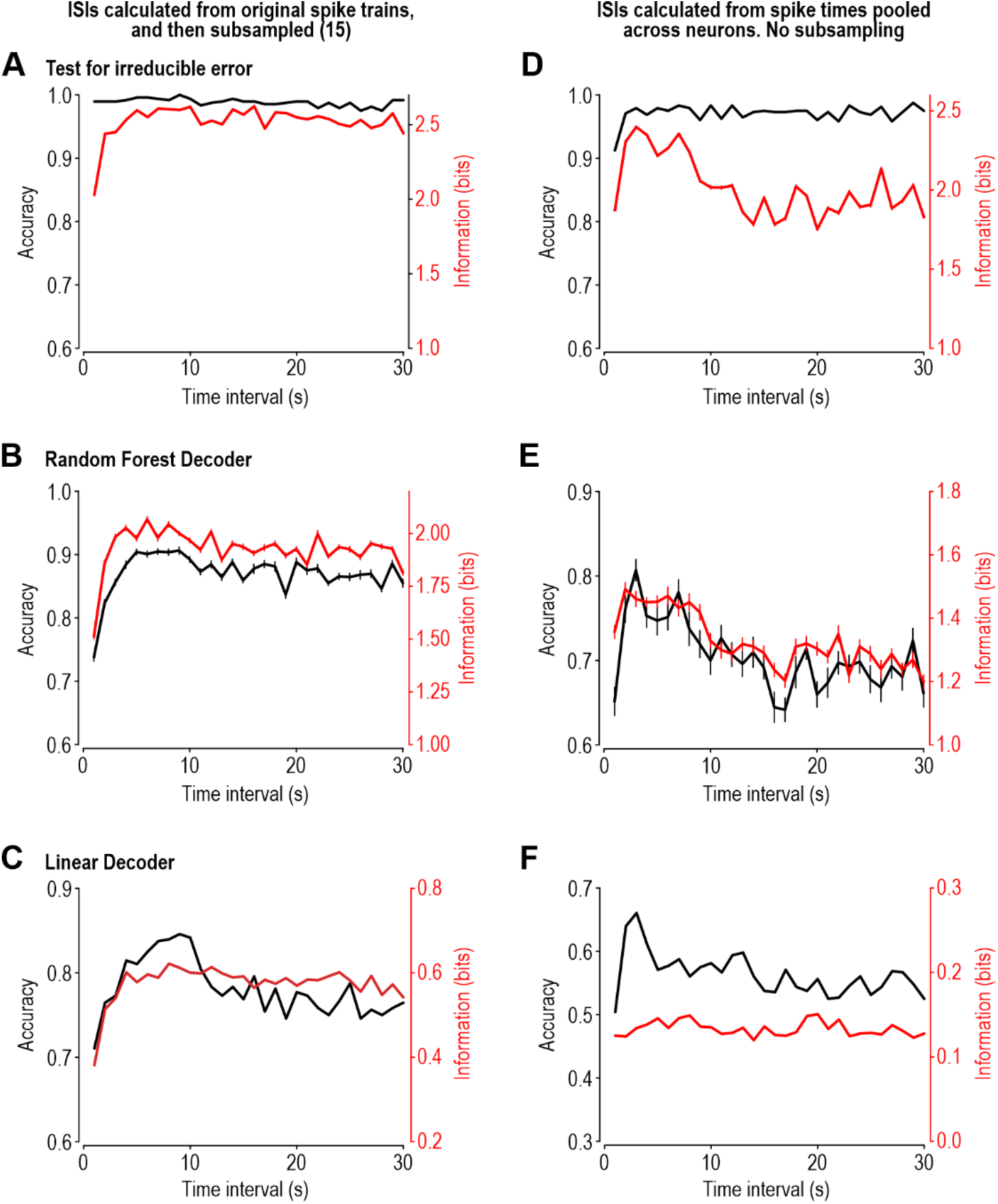
Comparison of decoders to assess discriminability of stimulus frequency from 15 ISIs subsampled from ISIs calculated from original spike trains (A-C) or from all ISIs calculated after spike times were pooled across neurons (D-F). **A.** Same as Fig. 4D pale but using an SVC but with the training set equal to the test set. **B.** Same as Fig. 4D pale but using a random forest decoder instead of an SVC. The train/test split (0.7/0.3) was repeated 20 times with different splits. **C.** Same as Fig. 4D pale but using a linear decoder instead of an SVC. The train/test split (0.7/0.3) was repeated 20 times with different splits. **D-F.** Same as A-C, respectively, but using ISIs calculated after spike times were pooled across neurons, like in Figure 4E.

**Figure S6.**
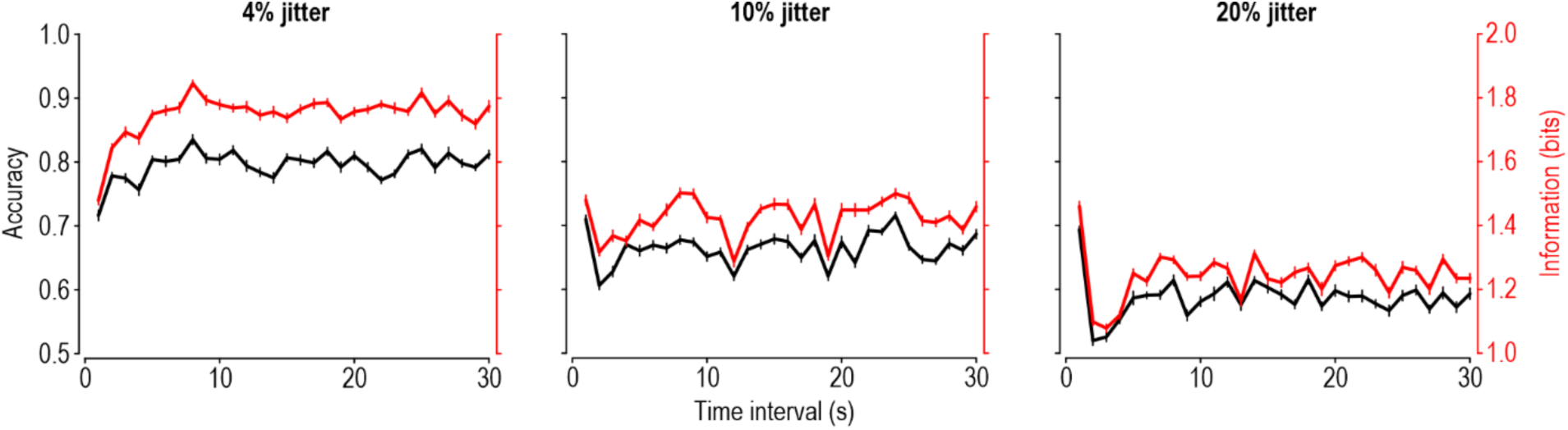
Extended analysis of effect of artificially altering phase-locking precision. Same as Fig. 5C pale (decoding of stimulus frequency from ISI distribution after jittering), but for spike times jittered by different amounts. Jitter spike times by <6% (left) did not diminish the gains in decidability afforded by adaptation, while strong jitter did (center and right).

## REFERENCES

1. Z. F. Mainen, T. J. Sejnowski, Reliability of spike timing in neocortical neurons. Science 268, 1503–1506 (1995).

2. P. Dayan, L. F. Abbott, Theoretical Neuroscience: Computational and Mathematical Modeling of Neural Systems (MIT Press, 2001).

3. A. L. Jacobs, et al., Ruling out and ruling in neural codes. Proc Natl Acad Sci USA 106, 5936–5941 (2009).

4. S. A. Prescott, S. Ratté, Pain processing by spinal microcircuits: Afferent Combinatorics. Curr Opin Neurobiol 22, 631–639 (2012).

5. S. A. Prescott, Q. Ma, Y. De Koninck, Normal and abnormal coding of somatosensory stimuli causing pain. Nat Neurosci 17, 183–191 (2014).

6. H. P. Saal, S. J. Bensmaia, Touch is a team effort: Interplay of submodalities in cutaneous sensibility. Trends Neurosci 37, 689–697 (2014).

7. K. Imaizumi, N. J. Priebe, T. O. Sharpee, S. W. Cheung, C. E. Schreiner, Encoding of temporal information by timing, rate, and place in cat auditory cortex. PLoS ONE 5 (2010).

8. E. L. Mackevicius, M. D. Best, H. P. Saal, S. J. Bensmaia, Millisecond precision spike timing shapes tactile perception. J Neurosci 32, 15309–15317 (2012).

9. S. Ratté, S. Hong, E. De Schutter, S. A. Prescott, Impact of neuronal properties on network coding: Roles of spike initiation dynamics and robust synchrony transfer. Neuron 78, 758– 772 (2013).

10. M. Lankarany, D. Al-Basha, S. Ratté, S. A. Prescott, Differentially synchronized spiking enables multiplexed neural coding. Proc Natl Acad Sci USA 116, 10097–10102 (2019).

11. J. W. Pillow, et al., Spatio-temporal correlations and visual signalling in a complete neuronal population. Nature 454, 995–999 (2008).

12. M. A. Harvey, H. P. Saal, J. F. Dammann, S. J. Bensmaia, Multiplexing stimulus information through rate and temporal codes in primate somatosensory cortex. PLoS Biol 11, e1001558 (2013).

13. H. P. Saal, M. A. Harvey, S. J. Bensmaia, Rate and timing of cortical responses driven by separate sensory channels. eLife 4, e10450 (2015).

14. M. A. Kamaleddin, et al., Physiological noise facilitates multiplexed coding of vibrotactile-like signals in somatosensory cortex. Proc Natl Acad Sci USA 119, e2118163119 (2022).

15. K. H. Long, J. D. Lieber, S. J. Bensmaia, Texture is encoded in precise temporal spiking patterns in primate somatosensory cortex. Nat Commun 13, 1311 (2022).

16. C. E. Shannon, A mathematical theory of communication. Bell System Technical Journal 27, 379–423 (1948).

17. A. Borst, F. E. Theunissen, Information theory and neural coding. Nat Neurosci 2, 947–957 (1999).

18. 18. D. J. C. MacKay, Information theory, Inference, and learning algorithms (Cambridge University Press, 2003).

19. 19. H. B. Barlow, Possible principles underlying the transformations of sensory messages. In Sensory Communication (MIT Press, 1961).

20. S. Laughlin, A simple coding procedure enhances a neuron’s information capacity. Z Naturforsch 36, 910–912 (1981).

21. N. Brenner, W. Bialek, R. de Ruyter van Steveninck, Adaptive rescaling maximizes information transmission. Neuron 26, 695–702 (2000).

22. A. L. Fairhall, G. D. Lewen, W. Bialek, R. R. de Ruyter van Steveninck, Efficiency and ambiguity in an adaptive neural code. Nature 412, 787–792 (2001).

23. T. O. Sharpee, et al., Adaptive filtering enhances information transmission in visual cortex. Nature 439, 936–942 (2006).

24. C. W. G. Clifford, et al., Visual adaptation: Neural, psychological and computational aspects. Vision Res 47, 3125–3131 (2007).

25. B. Wark, B. N. Lundstrom, A. Fairhall, Sensory adaptation. Curr Opin Neurobiol 17, 423– 429 (2007).

26. A. I. Weber, A. L. Fairhall, The role of adaptation in neural coding. Curr Opin Neurobiol 58, 135–140 (2019).

27. S. M. Smirnakis, M. J. Berry, D. K. Warland, W. Bialek, M. Meister, Adaptation of retinal processing to image contrast and spatial scale. Nature 386, 69–73 (1997).

28. T. Hosoya, S. A. Baccus, M. Meister, Dynamic predictive coding by the retina. Nature 436, 71–77 (2005).

29. M. Maravall, R. S. Petersen, A. L. Fairhall, E. Arabzadeh, M. E. Diamond, Shifts in coding properties and maintenance of information transmission during adaptation in barrel cortex. PLoS Biology 5, e19 (2007).

30. T. O. Sharpee, A. J. Calhoun, S. H. Chalasani, Information theory of adaptation in neurons, behavior, and mood. Curr Opin Neurobiol 25, 47–53 (2014).

31. S. G. Solomon, J. W. Peirce, N. T. Dhruv, P. Lennie, Profound contrast adaptation early in the visual pathway. Neuron 42, 155–162 (2004).

32. N. A. Lesica, et al., Adaptation to stimulus contrast and correlations during natural visual stimulation. Neuron 55, 479–491 (2007).

33. A. Benucci, A. B. Saleem, M. Carandini, Adaptation maintains population homeostasis in primary visual cortex. Nat Neurosci 16, 724–729 (2013).

34. K. W. Latimer, et al., Multiple timescales account for adaptive responses across sensory cortices. J Neurosci 39, 10019–10033 (2019).

35. M. Dipoppa, et al., Adaptation shapes the representational geometry in mouse V1 to efficiently encode the environment. bioRxiv (2024). 10.1101/2024.12.11.628035.

36. W. De Baene, E. Premereur, R. Vogels, Properties of shape tuning of macaque inferior temporal neurons examined using rapid serial visual presentation. J Neurophysiol 97, 2900– 2916 (2007).

37. Y. Shi, et al., Rapid, concerted switching of the neural code in inferotemporal cortex. bioRxiv (2023). 10.1101/2023.12.06.570341.

38. A. K. Goble, M. Hollins, Vibrotactile adaptation enhances amplitude discrimination. J Acoust Soc Am 93, 418–424 (1993).

39. A. K. Goble, M. Hollins, Vibrotactile adaptation enhances frequency discrimination. J Acoust Soc Am 96, 771–780 (1994).

40. I. Lampl, Y. Katz, Neuronal adaptation in the somatosensory system of rodents. Neuroscience 343, 66–76 (2017).

41. M. Adibi, I. Lampl, Sensory adaptation in the whisker-mediated tactile system: Physiology, theory, and function. Front Neurosci. 770011 **15** (2021).

42. M. A. Muniak, S. Ray, S. S. Hsiao, J. F. Dammann, S. J. Bensmaia, The neural coding of stimulus intensity: Linking the population response of mechanoreceptive afferents with psychophysical behavior. J Neurosci 27, 11687–11699 (2007).

43. S. Bensmaia, Tactile intensity and population codes. Behav Brain Res 190, 165–173 (2008).

44. A. I. Weber, et al., Spatial and temporal codes mediate the tactile perception of natural textures. Proc Natl Acad Sci USA 110, 17107–17112 (2013).

45. E. L. Graczyk, et al., The neural basis of perceived intensity in natural and artificial touch. Sci Transl Med 8, 362ra142 (2016).

46. H. P. Saal, X. Wang, S. J. Bensmaia, Importance of spike timing in touch: An analogy with hearing? Curr Opin Neurobiol 40, 142–149 (2016).

47. I. Birznieks, R. M. Vickery, Spike timing matters in novel neuronal code involved in vibrotactile frequency perception. Curr Biol 27, 1485–1490.e2 (2017).

48. G. Corniani, M. A. Casal, S. Panzeri, H. P. Saal, Population coding strategies in human tactile afferents. PLOS Comput Biol 18, e1010763 (2022).

49. H. P. Saal, B. P. Delhaye, B. C. Rayhaun, S. J. Bensmaia, Simulating tactile signals from the whole hand with millisecond precision. Proc Natl Acad Sci USA 114, E5693–E5702 (2017).

50. R. de Ruyter van Steveninck, W. Bialek, Real-time performance of a movement-sensitive neuron in the blowfly visual system: Coding and information transfer in short spike sequences. Proc R Soc Lond B Biol Sci 234, 379–414 (1988).

51. W. Bialek, F. Rieke, R. R. de Ruyter van Steveninck, D. Warland, Reading a neural code. Science 252, 1854–1857 (1991).

52. E. Zavitz, H.-H. Yu, E. G. Rowe, M. G. P. Rosa, N. S. Price, Rapid adaptation induces persistent biases in population codes for visual motion. J Neurosci 36, 4579–4590 (2016).

53. M. Snow, R. Coen-Cagli, O. Schwartz, Adaptation in the visual cortex: A case for probing neuronal populations with natural stimuli. F1000Research 6, 1246 (2017).

54. 54. T. Hastie, R. Tibshirani, J. Friedman, The Elements of Statistical Learning, second edition: Data Mining, Inference, and prediction (Springer, 2009).

55. P. Sterling, S. Laughlin, Principles of Neural Design (The MIT Press, 2017).

56. 56. J. V. Stone, Principles of Neural Information Theory: Computational Neuroscience and Metabolic Efficiency (Sebtel Press, 2018).

57. E. D. Adrian, Y. Zotterman, The impulses produced by sensory nerve-endings. J Physiol 61, 151–171 (1926).

58. D. R. Ollerenshaw, H. J. V. Zheng, D. C. Millard, Q. Wang, G. B. Stanley, The adaptive trade-off between detection and discrimination in cortical representations and behavior. Neuron 81, 1152–1164 (2014).

59. B. N. Lundstrom, M. H. Higgs, W. J. Spain, A. L. Fairhall, Fractional differentiation by neocortical pyramidal neurons. Nat Neurosci 11, 1335–1342 (2008).

60. B. Wark, A. Fairhall, F. Rieke, Timescales of inference in visual adaptation. Neuron 61, 750–761 (2009).

61. M. Tommerdahl, et al., Human vibrotactile frequency discriminative capacity after adaptation to 25 Hz or 200 Hz stimulation. Brain Res 1057, 1–9 (2005).

62. J. A. Pruszynski, R. S. Johansson, Edge-orientation processing in first-order tactile neurons. Nat Neurosci 17, 1404–1409 (2014).

63. L. Duong, C. Bredenberg, D. Heeger, E. Simoncelli, Adaptive coding efficiency in recurrent cortical circuits via gain control. arXiv (2023). 10.48550/arXiv.2305.19869.

64. A. I. Weber, K. Krishnamurthy, A. L. Fairhall, Coding principles in adaptation. Annu Rev Vis Sci 5, 427–449 (2019).

65. J. Mathews, A. (Jaelyn) Chang, L. Devlin, M. Levin, Cellular signaling pathways as plastic, proto-cognitive systems: Implications for biomedicine. Patterns 4, 100737 (2023).

66. J. Thorson, M. Biederman-Thorson, Distributed relaxation processes in sensory adaptation. Science 183, 161–172 (1974).

67. B. N. Lundstrom, Modeling multiple time scale firing rate adaptation in a neural network of local field potentials. J Comput Neurosci 38, 189–202 (2014).

68. T. Sharpee, N. C. Rust, W. Bialek, Analyzing neural responses to natural signals: Maximally informative dimensions. Neural Comput 16, 223–250 (2004).

69. K. W. Latimer, A. L. Fairhall, Capturing multiple timescales of adaptation to second-order statistics with generalized linear models: Gain scaling and fractional differentiation. Front Syst Neurosci 14, 60 (2020).

70. L. T. McIntosh, N. Maheswaranathan, A. Nayebi, S. Ganguli, S. A. Baccus, Deep Learning Models of the Retinal Response to Natural Scenes. Adv Neural Inf Process Syst 29, 1369– 1377 (2016).

71. D. Beniaguev, I. Segev, M. London, Single cortical neurons as deep artificial neural networks. Neuron 109, 2727–2739 (2021).

72. P. McMillen, M. Levin, Collective intelligence: A unifying concept for integrating biology across scales and substrates. Commun Biol 7, 378 (2024).

73. P. C. Schwindt, W. J. Spain, R. C. Foehring, M. C. Chubb, W. E. Crill, Slow conductances in neurons from cat sensorimotor cortex in vitro and their role in slow excitability changes. J Neurophysiol 59, 450–467 (1988).

74. K. J. Kim, F. Rieke, Temporal contrast adaptation in the input and output signals of salamander retinal ganglion cells. J Neurosci 21, 287–299 (2001).

75. X.-J. Wang, Y. Liu, M. V. Sanchez-Vives, D. A. McCormick, Adaptation and temporal decorrelation by single neurons in the primary visual cortex. J Neurophysiol 89, 3279–3293 (2003).

76. H. J. Abel, J. C. F. Lee, J. C. Callaway, R. C. Foehring, Relationships between intracellular calcium and afterhyperpolarizations in neocortical pyramidal neurons. J Neurophysiol 91, 324–335 (2004).

77. S. A. Prescott, T. J. Sejnowski, Spike-rate coding and spike-time coding are affected oppositely by different adaptation mechanisms. J Neurosci 28, 13649–13661 (2008).

78. L. Paninski, J. Pillow, J. Lewi, Statistical models for neural encoding, decoding, and optimal stimulus design. Prog Brain Res 165, 493–507 (2007).

79. L. R. Manfredi, et al., Natural scenes in tactile texture. J Neurophysiol 111, 1792–1802 (2014).

80. B. P. Delhaye, H. P. Saal, S. J. Bensmaia, Key considerations in designing a somatosensory neuroprosthesis. J Physiol (Paris) 110, 402–408 (2016).

81. L. Medlock, et al., Encoding of vibrotactile stimuli by mechanoreceptors in rodent glabrous skin. J Neurosci 44, e1252242024 (2024).

